# Development of a metabolic signature of post-weaning diarrhoea in pigs

**DOI:** 10.1101/2025.11.17.688604

**Authors:** Joanna C. Wolthuis, Stefanía Magnúsdóttir, Edwin Stigter, Yuen Fung Tang, Judith Jans, Myrthe Gilbert, Bart van der Hee, Pim Langhout, Walter Gerrits, Arie Kies, Jeroen de Ridder, Saskia van Mil

## Abstract

In the agricultural sector, antibiotics have been used to improve swine growth performance. This application is banned nowadays, due to increased risk of antibiotic resistance. In piglets this results in a higher prevalence of post-weaning diarrhoea, deteriorating both animal health and performance. Our goal was to find a metabolite signature separating piglets with low faecal consistency score (FCS) from piglets with normal faecal consistency and determine which pathways were enriched in this signature.

By using direct infusion mass spectrometry on blood spots, we built machine learning (ML) models that aimed to differentiate between low and normal FCS. To test the general predictive capability of these models, we applied a Leave-One-Country-Out (LOCO) strategy for cross validation. Our second approach after LOCO was finding the optimal number of features to include in a feature-reduced model. To determine the order in which features were to be eliminated, we ranked them based on a combination of t-test and fold-change significance scores. Enrichment analysis using mummichog was used to gain insights into the final signature set of m/z values found using this ranking and ML models.

Models trained both using all countries and leaving out specific countries from training showed a limited ability to predict FCS category. Furthermore, the LOCO results were mixed, with some countries showing a predictive signal present in the data, but others with predictive capability that was no better than random.

Signature analysis using t-test and fold-change results did not result in any KEGG pathways that were enriched in this signature as compared to random.

Using these methods, we could predict the FCS category to a limited degree.

Although no common signature for low faecal consistency could be determined using this method, given that some countries showed more reliable LOCO results, further analysis into specifically the samples from those countries could be a valuable next step. No metabolite signature enriched in changes to KEGG pathways was found in the data using the combined t-test and fold-change analysis ranking method. We have shared the data with the community through the *MetaboLights* repository.

## Introduction

Consistent use of in-feed antibiotics to promote animal growth has been forbidden in many countries due to increased antibiotic resistance in the population, causing major concern for both animal and human health and leading to subsequent changes in public opinions towards these antibiotic growth promoters (AGPs) (Rahman, Fliss, and Biron 2022). Unfortunately, discontinued use of these AGPs can lead to health problems in the previously chronically treated animals, such as failure to thrive and subsequent higher mortality.

In pigs, this decrease in health is connected to less solid faecal consistency in early life, specifically post-weaning, defined as ‘post-weaning diarrhoea’ (PWD). At 3-4 weeks of age, piglets are weaned from their mother and their diet is changed from sow’s milk to starter feed, which is usually presented in a solid form. Post-weaning diarrhoea is a multifactorial disease, related to this dietary change in an intestine that is still developing (Gilbert et al. 2018) Alongside microbial imbalances and infection by bacteria and viruses, this may result in PWD, but the exact cause(s) remain(s) unclear (Gilbert et al. 2018).

Worldwide, PWD is a large problem causing high mortality in the farming industry (Nabuurs, Van Zijderveld, and De Leeuw 1993; Svensmark et al. 1989; Amezcua et al. 2002). Unfortunately, due to the fact that the housed pig genetic strains differ per country, alongside variation in animal housing and management and differences in native pathogens, it remains challenging to uniformly characterise this disease as physiological causes may differ.

To further prevent and potentially treat disease without antibiotic use, it would be highly beneficial to gain more insight into similarities and differences of this phenotype between and across countries. In chickens, we achieved this using metabolomic analysis to determine a metabolic signature(Wolthuis et al. 2023).

Metabolomics technology is flexible and allows for health classification, monitoring and potentially future disease intervention. Direct infusion mass spectrometry (DI-MS) is capable of mapping the metabolic profile from just a single drop of blood. This high-throughput method has an untargeted nature, which allows for minimization of molecule selection bias. In chickens, DI-MS was used to generate data from blood samples of healthy and unhealthy chickens, and the resulting data was processed using our software solutions *MetaboShiny* and *MetaDBparse* to normalise, statistically analyse and classify the data through machine learning(Wolthuis et al. 2020, 2023).

Here, we aimed to derive a metabolic signature of PWD that is shared across countries, hopefully thereby minimising the impact of differences in environment, feed and genetic strain. Such a signature would be an important source of insights on the biological processes connected to PWD, or to predict PWD at an early stage. From those insights, potential preventative and treatment plans may be developed in the future. Blood samples of piglets with and without PWD were collected from 24 farms from four countries across three continents. We applied both univariate statistics and machine learning to investigate the presence of a metabolic signature underlying PWD in piglets.

## Results

### Metabolic profiling from blood

An overview of our methodology is presented in *Figure 1*. On-site experts selected 330 pigs with low to high faecal consistency ranging from 1 to 5 based on a provided chart (*Figure 1a-b, Table S1*). Supervised by local veterinarians or technicians, pigs were selected and sampled from 24 farms in Australia, the Netherlands, the USA and Brazil. The blood samples were taken on blood spot cards and dried. Alongside the recorded Faecal Consistency Score (FCS), body weight, sex, feed, housing type (open, closed, ventilated), environmental temperature and genetic strain of each animal was registered in a provided spreadsheet.

**Figure 1.**
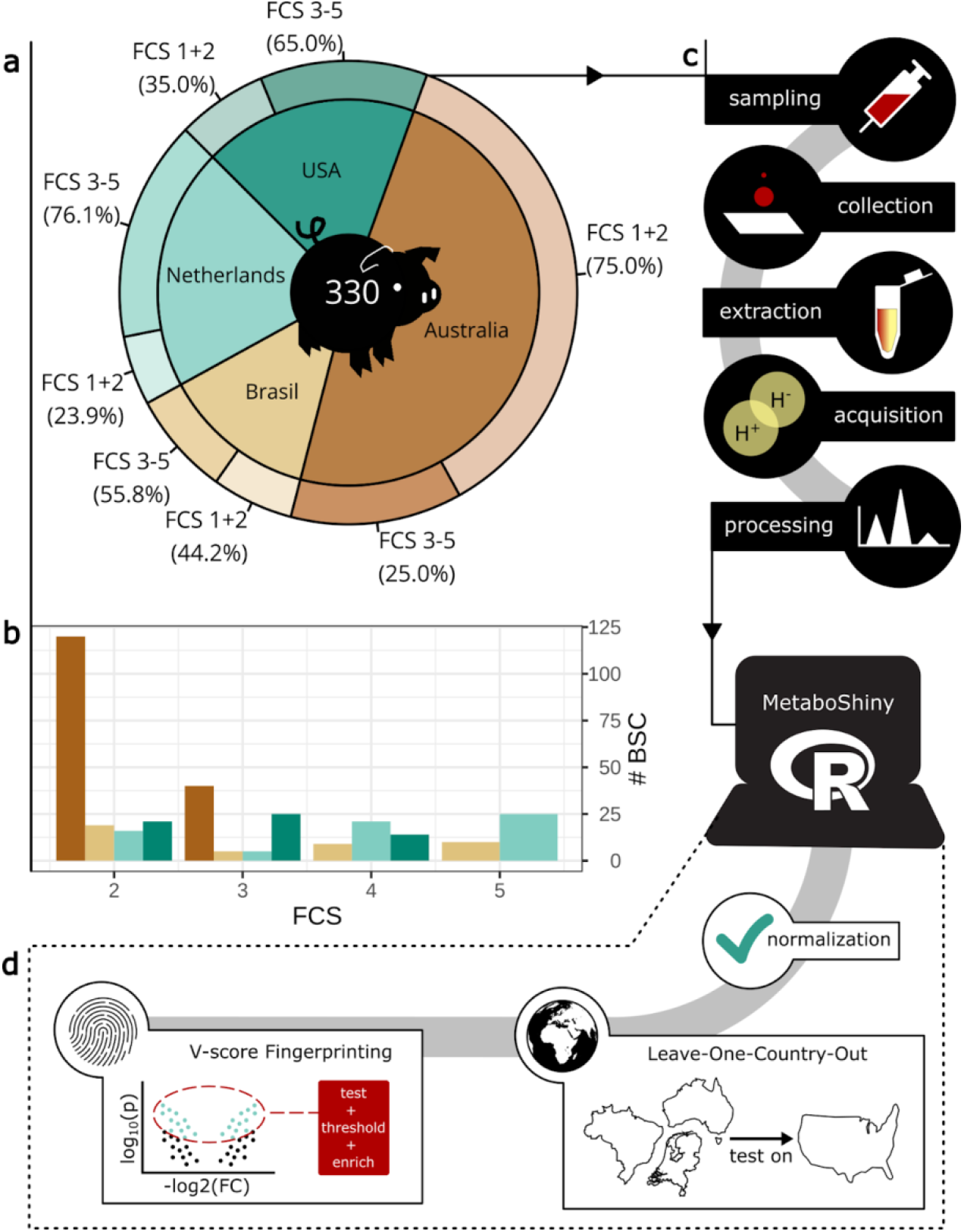
Worldwide pig sample collection and analysis of faecal consistency score. **(a)** Overview of the ratio of samples collected from the four participating countries. **(b)** Faecal Consistency Score (FCS) distribution of samples per country. **(c)** Blood samples were collected on blood spot cards, subsequently extracted and signal acquired through mass spectrometry. Bioinformatic processing prepared the data for analysis through MetaboShiny. **(d)** Within MetaboShiny, two main analyses were performed. Leave-One-Country-Out where machine learning was used to test the predictive performance on a separate country, and V-score signature searching which combines two univariate tests to rank mass/charge values, to subsequently narrow down a predictive signature consisting of the highest-ranking features.

To analyse the samples, high-resolution chip-based electrospray mass spectrometry was performed in-house, and the subsequent spectra were processed using the pipeline described by (de Sain-van der Velden et al. 2017). Alignment, summing and binning of the data was performed in accordance with (Wolthuis et al. 2023) to prepare the data for MetaboShiny analysis (*Figure 1c*).

Metadata-wise, the piglets were divided into two groups, a group containing FCS 1 - 2 (low-FCS), and a second group containing FCS 3 - 5 (high-FCS) to enable two-class machine learning model construction. Groups of FCS 1-3 and FCS 4-5 were not chosen as not all countries could provide samples with FCS > 3.

Within MetaboShiny, data was normalised and batch corrected with *waveICA* (Deng et al. 2021). Initially, batches show separation in PCA, and post-processing this separation is decreased, allowing for analysis with reduced influence of batch effects (*Figure S1*).

### Leave-One-Country-Out analysis shows no difference between FCS categories

We aimed to find a metabolite signature separating the low-FCS group from the high-FCS group, preferably one that is applicable worldwide (*Figure 1d*). To test the effectiveness of our model on other countries, we approached the problem using Leave-One-Country-Out (LOCO) analysis, where cross-validation was applied by excluding one country from training and using it as the testing group.

These models used Random Forest classification without limiting the features (m/z values) used. As a baseline, we first built a model using all countries in each training and testing fold, the ‘multi-country’ model, which can be used to evaluate the other models. The results of the subsequent LOCO analysis are shown in *Figure 2* through both AUPRC (Area under the Precision-Recall Curve) and AUROC (Area Under the Receiver Operating Characteristic), which both would equal 1 in a maximally predictive model, whilst significantly differing from the shuffled (red) model.

**Figure 2.**
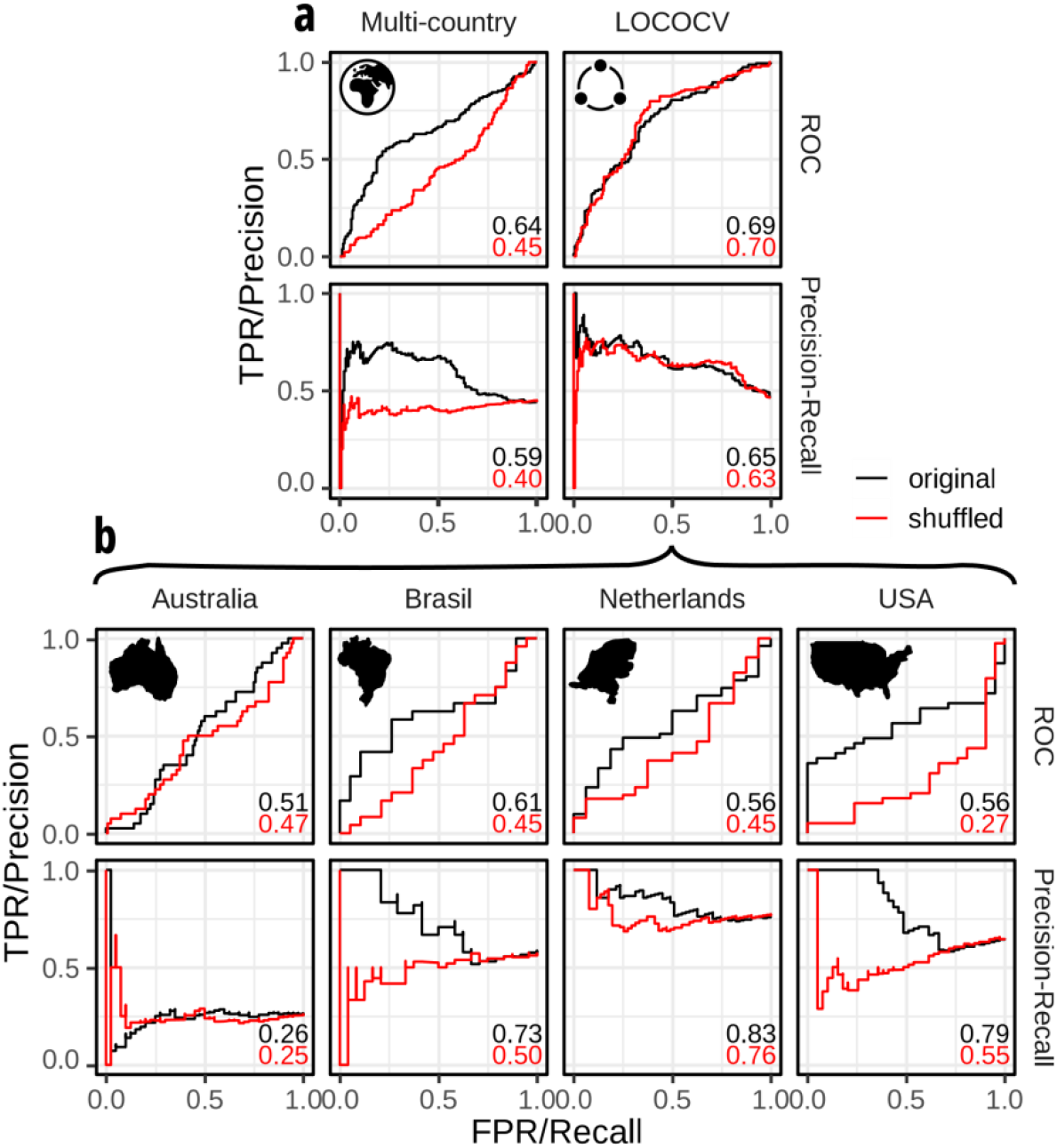
Leave-One-Country-Out-Cross-Validation (LOCOCV) and multi-country model Receiver Operator Characteristic (ROC) and Precision-Recall (PR) curves. Performance in AUPRC and AUROC of original (black) and shuffled label (red) models is noted in lower right. **(a)** Multi-country model uses all countries simultaneously in 10-fold cross-validation. LOCOCV model testing folds are composed of individual countries. The LOCOCV curves summarise per-country performance. **(b)** Performance of separate LOCO models. The mentioned country is the used testing set, the other three countries are used as training.

This multi-country model performed better than random with an Area Under the Receiver Operator characteristic Curve (AUROC) of 0.64 and Area Under the Precision-Recall Curve (AUPRC) of 0.59. Next, we built the Leave-One-Country-Out-Cross-Validation (LOCOCV) model, and evaluated it by aggregating the test sets (the left-out countries). Altogether, the LOCOCV model performed worse than the multi-country model, as the AUROC is similar (0.69), the AUPRC is better (0.65), but we see no difference between a negative-control model using non-shuffled versus shuffled labels.

We next analysed the separate folds of the LOCOCV to establish the similarity of a specific country’s FCS signature to the other training countries. This is ideally done using the AUPRC due to the class imbalances present in the dataset. Brazil (AUROC/AUPRC: 0.61/0.73) and USA (0.56/0.79) perform best, showing large differences between the negative control and experiment (ΔAUPRC Brazil: 0.23, USA:0.49), followed by the Netherlands (AUROC/AUPRC: 0.56/0.83) and Australia (AUROC/AUPRC: 0.51/0.26).

### Signature from V-score feature selection is not enriched in pathways

We continued analysis using the multi-country model, and in order to investigate what metabolic signature underlies the predictive value of this model, we proceeded aiming to find the maximum number of m/z values of value for predicting low- or high-FCS samples. First, the data was split into 80/20 fractions, of which the 80% fraction was used for signature discovery. We re-used the 10 folds defined in the multi-country model, and per fold we performed a t-test and fold-change analysis in the majority fraction.

Our aim was to rank the features based on our previously published ‘V-score’ metric, a combination of transformed t-test and fold-change outcomes (Wolthuis et al. 2023). We calculated the V-score for each m/z value by combining the results of these univariate tests.

This ranking was followed by building a Random Forest model with a decreasing number of m/z values, 50 at a time, starting with the highest V-score-ranked features. Per step of leaving out these m/z values, we also included a negative control for the ranking, which removes 50 random m/z values (*Figure 3a*, blue line), and a negative control for the class labels with shuffled labels (*Figure 3a*, red line). This leads to a series of models with decreasing numbers of features. We applied these models to the minority fold and calculated the AUROC *(Figure 3a)*.

**Figure 3.**
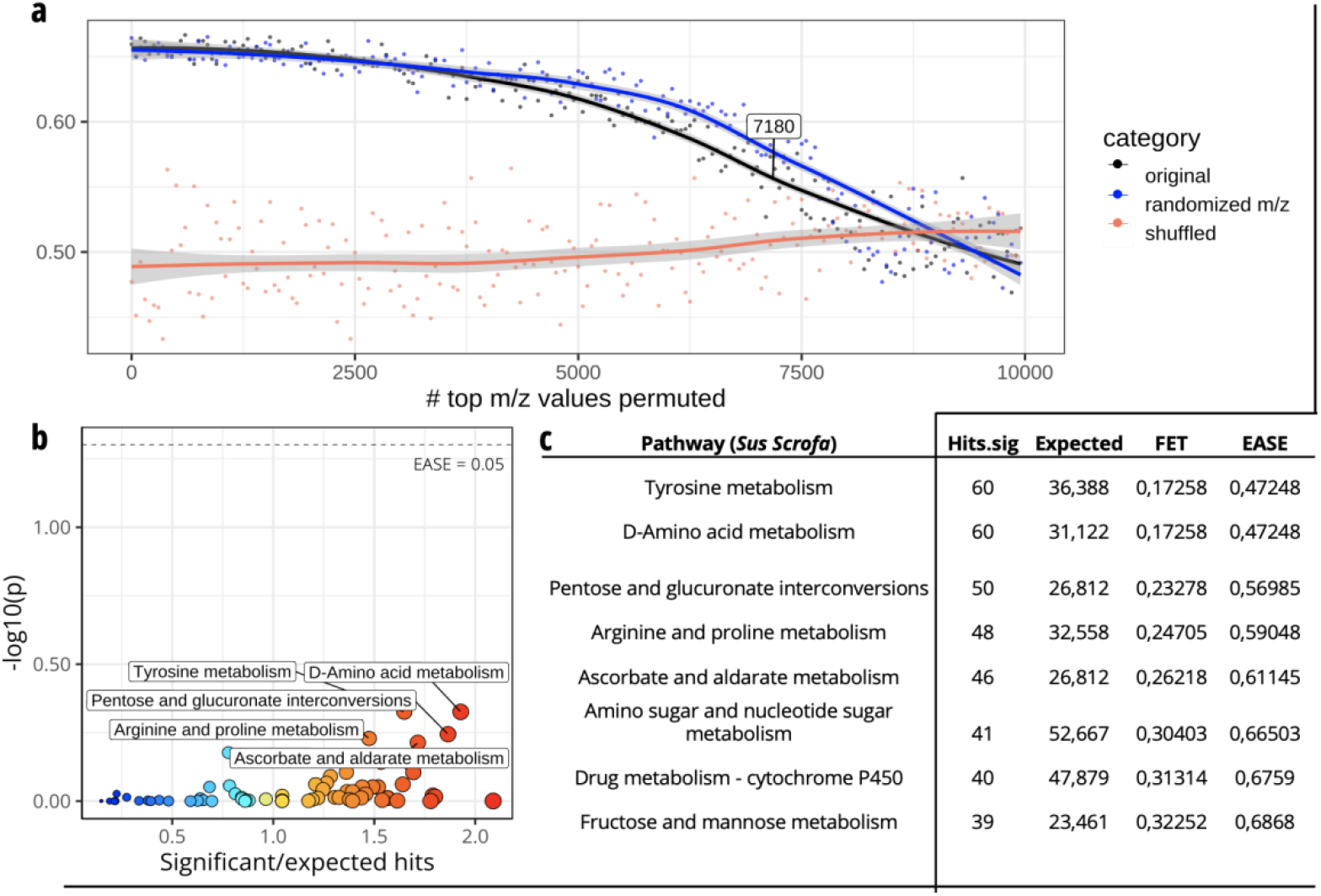
Signature discovery based on m/z V-score ranking. **(a)** Predictive value curves for the experimental model (black line, removing ranked features), negative control 1 (blue line, removing random features) and negative control 2 (red line, removing ranked features but with random FCS labels). **(b)** No pathways are significantly enriched at EASE < 0.05 (dashed line) for the *Sus scrofa* KEGG database. **(c)** Top 10 enriched pathways in 7180-m/z top V-score ranked signature. FET=Fischer’s Exact Test. EASE: conservative version of FET used for our analyses.

We observed that the experimental and negative control curves did not differ majorly, with only a slight dip present in the ranked-removal curve with an elbow at 7180 m/z values. On the 20% of the data left out at the advent of the experiment, we calculated the V-score and performed enrichment analysis on the top 7180 m/z values using mummichog and the *Sus scrofa* database. No significant pathways were discovered at EASE (conservative version of Fisher’s Exact test) < 0.05.

## Methods

### Sample acquisition

Four drops of blood were collected from each of 330 pigs from 4 countries on “Whatman™ 903 Protein Saver Cards” blood spot cards (BSCs) (Sigma-Aldrich Merck KGaA, Darmstadt, Germany). Brazil supplied 43 BSCs from 10 farms, the United States of America (USA) provided 60 BSCs from 6 farms, the Netherlands supplied 67 BSCs from four farms, and Australia provided 160 BSCs from four farms (*Figure 1*). These BSCs were sealed in plastic bags alongside humidity monitoring cards and desiccant and shipped to the UMC Utrecht, and stored at -80C until further processing. Metadata was collected for each animal, on faecal consistency score based on the guides visualised in *Tables S1* and *S2*, feed type, age, body weight, genetic strain, housing type and environmental temperature.

### Data collection

Using our in-house pipeline for DI-MS (de Sain-van der Velden et al. 2017), we extracted three 3mm punches from the blood spot card for three technical replicates. Extraction was performed in an ultrasonic bath using acetonitrile, formic acid and internal standards.

Subsequently, the resulting extract was filtered and measured in DI-MS analysis in both positive and negative mode using a Thermo Scientific Q-Exactive HF high-resolution mass spectrometer equipped with an Advion Triversa Nanomate. Positive and negative signals were collected for 3 minutes and 1.5 minutes respectively. Resulting data was stored in .*raw* Thermo format.

### Processing and normalisation

The Thermo RAW files were formatted to .*mzML* using ThermoFileReader and further processing such as splitting into positive and negative mode, alignment and summing of replicates, peak gathering and binning were performed according to (Wolthuis et al. 2023). The used scripts can be found on the Github *joannawolthuis/MassChecker* repository. A m/z value was only retained if it was found present in more than 20% of the samples in the dataset (Bijlsma et al. 2006). Autoscaling, quantile normalisation and batch correction using *waveICA* were used within MetaboShiny normalisation settings (Deng et al. 2021).

### Feature selection and data distribution

The data was split in two sections. The first 80% was used for feature selection-based machine learning, within which 10 folds were defined for cross-validation purposes. The other 20% was retained for enrichment analysis on the signature found in the other fraction of the data.

Feature selection was based on the ‘V-score’, a multiplication of -log10(t-test p-value) and the log2(fold-change) and performed in accordance with (Wolthuis et al. 2023). For Leave-One-Country-Out analysis, data was separated purely based on country of origin.

### Signature determination

With half of the samples, we defined 10 folds. In each fold, we calculate the V-score and train a random forest model, which is then tested on the held-out data of that fold. This is done for each of the 10 folds, and predictive performance is recorded. This is done while shrinking the feature set by 50 m/z at a time, and removing the top V-score ranking features first. With the resulting AUC values a curve is drawn. This experiment is repeated with two negative controls, a shuffled-(FCS category)-label negative control, and a model that removes 50 random features. The signature is determined by finding the point in the result curve with the maximum difference (elbow point) of the line fit representing the V-score ranked models.

### Enrichment

Enrichment analysis was performed using mummichog within MetaboShiny (Wolthuis et al. 2020; Li et al. 2013). We used the ‘*Sus scrofa’* (KEGG: ssc01100) and ‘microbial metabolism in diverse environments’ (KEGG: map01120) databases as metabolite and pathway source (Kanehisa and Goto 2000). We used the adducts described in table S3.

Additionally, to reduce the considered possible pathways and thus increase statistical power, a filtering step was performed based on the essential amino acids in pigs: histidine, isoleucine, leucine, lysine, methionine, phenylalanine, threonine, tryptophan, and valine (Rezaei et al. 2013). This leads to the exclusion of KEGG pathways ‘*valine, leucine and isoleucine biosynthesis’* and *‘phenylalanine, tyrosine and tryptophan biosynthesis*’. We marked all metabolites up to the signature threshold of 7180 as significant, and pathways were considered significantly enriched if the EASE p-value was below or equal to 0.05. This approach was limited to strictly essential amino acids to ensure biosynthesis of these amino acids is disabled and thus these pathways are impossible.

### Visualisation

Plotting was performed within MetaboShiny using *ggplot2, ggrepel*, and *moonBook* and their dependencies (Wolthuis et al. 2020; Wickham 2011). To visualise the signature determination based on V-score, we fit a line using the *ggplot2 geom_smooth* function to the measured AUC values for the respective groups. We determined the signature threshold in *Figure 3a* by finding the sharpest curve point of the line representing the ranked m/z values in their original order (black line).

## Discussion and conclusion

We set out to gain insight into the metabolic changes connected to post-weaning diarrhoea in pigs, to potentially elucidate causal and consequential mechanisms of this phenotype. To do so, we built models using the blood metabolome in pigs to predict FCS categories. These models, if found to be significantly predictive, could be used to subsequently gain insights in the metabolic changes underlying a high or low FCS, if a m/z signature was found.

Our ‘Leave-One-Country-Out’ (LOCO) experiment was designed to evaluate the potential of our models to predict FCS category in a country unknown to the training dataset (Figure 2). The LOCO model underperformed compared to the multi-country model, suggesting that the underlying differences between countries are substantially large. This could be due to different causes of the low-FCS phenomenon between countries, differences in environment, feed composition, and genetic background of the animals (Park, Min, and Oh 2017). Further research may map these differences and investigate potential methods to use this part of the metadata to build a model using more non-metabolomic features.

On the multi-country level, we next aimed to determine a m/z signature connected to the difference between the low- and high-FCS groups (Figure 3). We combined the t-test and fold-change univariate analyses into a multiplicative score called the ‘V-score’ that we have previously applied to chicken data (Wolthuis et al. 2023). The goal of this ranking prior to model construction was to make sure that the used features were more likely to be interpretable on an individual m/z basis.

We divided the data into 10 folds, and in each fold, we calculated the V-score, ranked the features, and then built a model that was tested on the left-out data. This model used a decreasing number of m/z features - the top V-score ranking m/z values were removed to investigate the effect of removal on the model predictive value. The difference between our negative control removing random features, and our experiment removing highly ranked features, was low, and the resulting m/z signature showed no significant enrichment in *Sus scrofa* KEGG pathways (*Figure 3b-c)*. This indicates that the m/z values selected through this analysis likely do not capture any biologically significant differences separating the low and high-FCS groups.

We investigated the metabolic changes behind the differences between high- and low-FCS pigs to find causes and/or consequences of this phenotype on blood level. Unfortunately, despite this methodology resulting in a clear metabolic signature in poultry (Wolthuis et al. 2022), we were not able to successfully identify a metabolic signature in pigs. Potential causes of this are large genetic, feed- and environment-based differences between countries and/or farms, complicating extraction of a clear underlying metabolic signal. Another potential cause may be the chosen two-group separation. Although 1-2 and 3-5 were chosen to enable analysis of samples from all countries with an optimal class balance, it may be worth considering comparing the FCS 1-3 with FCS 4-5 groups in future studies.

We hope that by sharing our data with the community, this dataset, rich in metadata, can be used to continue this line of research or investigate the blood metabolome in the context of sex, genetic line, feed, housing and body weight. This addition of 330 pigs to the *MetaboLights* database will enrich the existing database of samples and facilitate future studies in pigs.

## Data availability

The data used in this study is available in *MetaboLights* study MTBLS8937.

## Ethical statements

Animals were cared for in accordance with all applicable institutional, national and/or international guidelines.

## Funding contributions

This study was supported by the Applied Science Division (STW) of the Dutch Organisation for Scientific Research (Nederlandse Organisatie voor Wetenschappelijk Onderzoek, NWO) alongside DSM Nutritional Products. Jeroen de Ridder is supported by the Vidi NWO Fellowship (614.072.715).

