## Supplemental Materials for "Development of a metabolic signature of post-weaning diarrhoea in pigs"

*Table S1: Faecal consistency scoring*

| FCS | Description | Image |
|---|---|---|
| 1 | Hard (no diarrhoea) | N/A |
| 2 | Normal (no diarrhoea) | 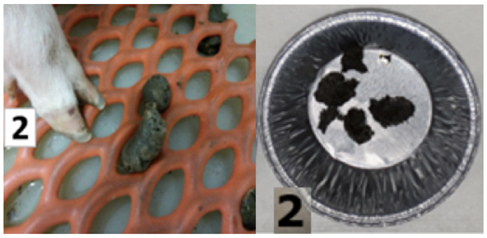 |
| 3 | Soft (diarrhoea) | N/A |
| 4 | Paste (diarrhoea) | 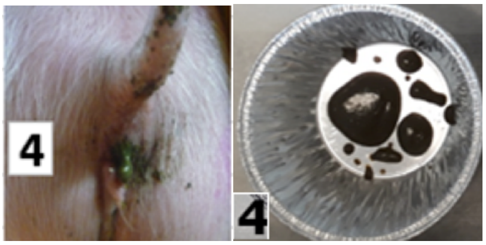 |
| 5 | Liquid (diarrhoea) | 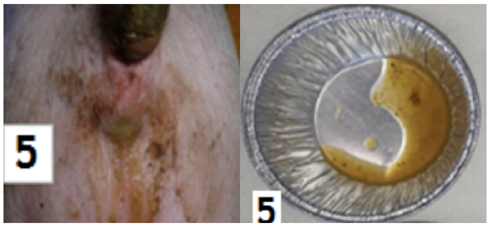 |

*Table S2: Faecal consistency scoring on grid*

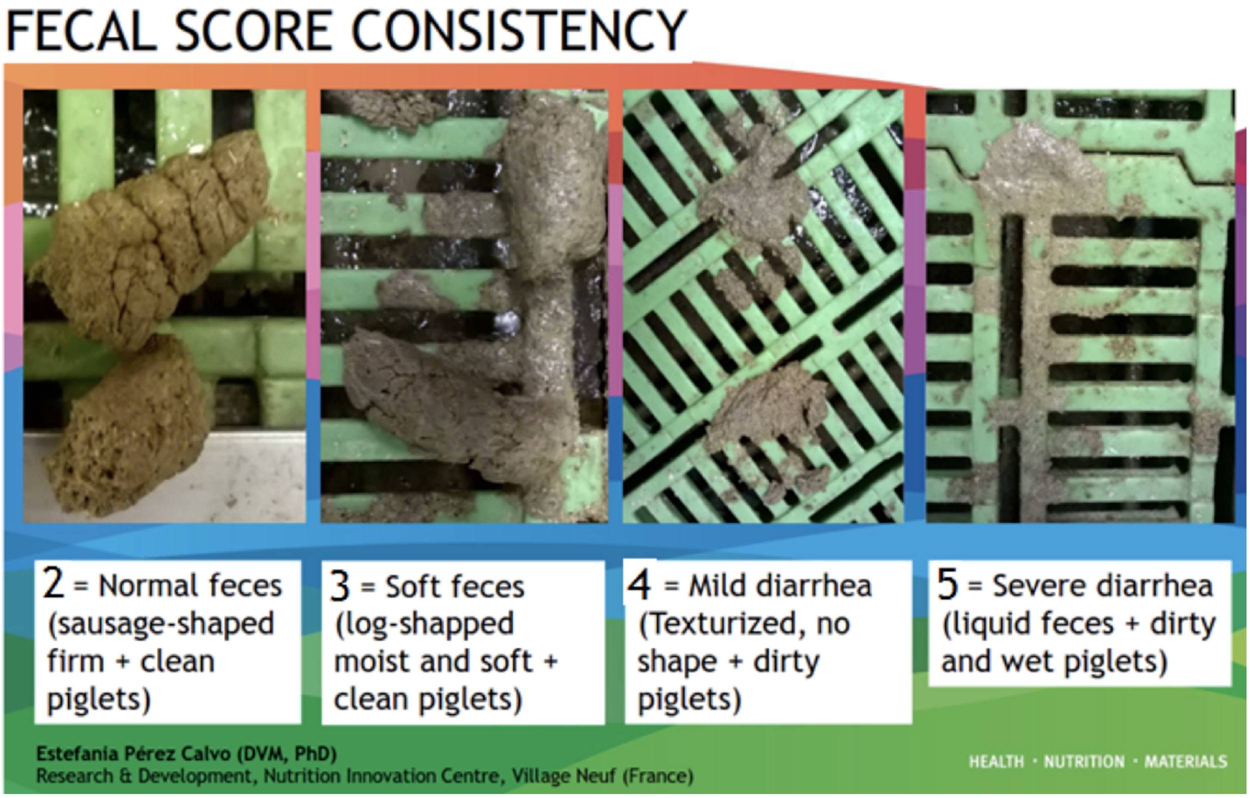

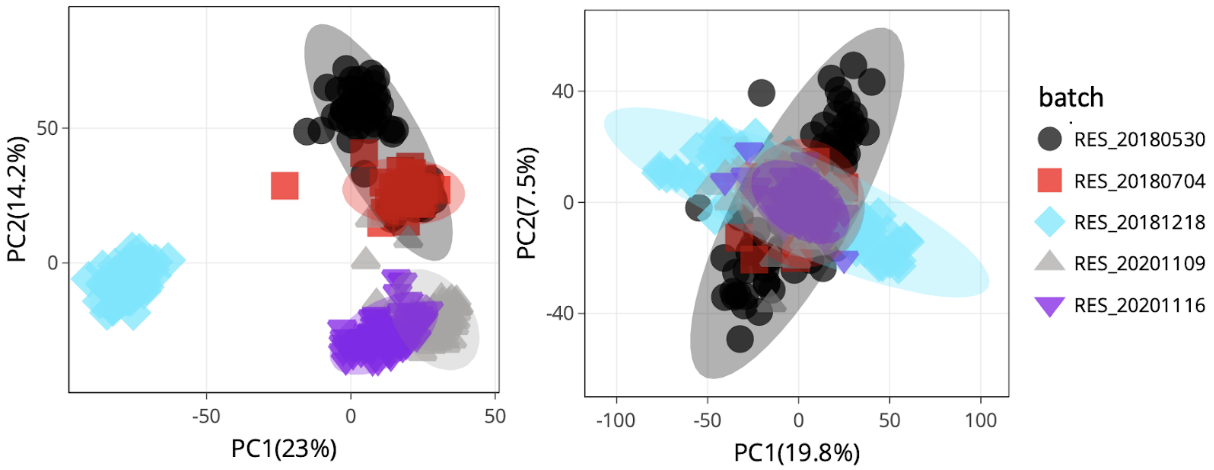


**Figure S1:** PCA of data pre- and post-batch correction. Shape and colour indicate day-of-run batch.

### *Table S3: Adduct table*

| Name | Ion_mode | Charge | AddEx | RemEx | Nelec | Rule |
|---|---|---|---|---|---|---|
| [M-H+Cl]2- | negative | -2 | Cl1 | H1 | 2 | Ndon>0 AND Nch=0 |
| [M-H+FA]1- | negative | -1 | H2C1O2 | H1 | 1 | Ndon>0 AND Nch=0 |
| [M+H]1+ | positive | 1 | H1 | | -1 | Nacc>0 AND Nch=0 |
| [M+K]1+ | positive | 1 | K1 | | -1 | Nacc>0 AND Nch=0 |
| [M+Na]1+ | positive | 1 | Na1 | | -1 | Nacc>0 AND Nch=0 |
| [M+NH4]1+ | positive | 1 | N1H4 | | -1 | Nacc>0 AND Nch=0 |
| [M1+.]1+ | positive | 1 | | | 0 | Nch=1 |
| [M-H]1- | negative | -1 | | H1 | 1 | Ndon>0 AND Nch=0 |
| [M+Cl]1- | negative | -1 | Cl1 | | 1 | Nacc>0 AND Nch=0 |
| [M+NaCl+H]1+ | positive | 1 | Na1Cl1H1 | | -1 | Nacc>0 AND Nch=0 |
| [M+(NaCl)2+H]1+ | positive | 1 | Na2Cl2H1 | | -1 | Nacc>0 AND Nch=0 |
| [M+(NaCl)3+H]1+ | positive | 1 | Na3Cl3H1 | | -1 | Ndon>0 AND Nch=0 |
| [M+(NaCl)4+H]1+ | positive | 1 | Na4Cl4H1 | | -1 | Nacc>0 AND Nch=0 |
| [M+(NaCl)5+H]1+ | positive | 1 | Na5Cl5H1 | | -1 | Nacc>0 AND Nch=0 |
| [M+NaCl-H]1- | negative | -1 | Na1Cl1 | H1 | 1 | Ndon>0 AND Nch=0 |
| [M+(NaCl)2-H]1- | negative | -1 | Na2Cl2 | H1 | 1 | Ndon>0 AND Nch=0 |
| [M+(NaCl)3-H]1- | negative | -1 | Na3Cl3 | H1 | 1 | Ndon>0 AND Nch=0 |
| [M+(NaCl)4-H]1- | negative | -1 | Na4Cl4 | H1 | 1 | Ndon>0 AND Nch=0 |
| [M+(NaCl)5-H]1- | negative | -1 | Na5Cl5 | H1 | 1 | Ndon>0 AND Nch=0 |

*Addex =  Added atoms

*RemEx = removed atoms

*Nelec = electron amount
